# Sex-specific genetic effects across biomarkers

**DOI:** 10.1101/837021

**Authors:** Emily Flynn, Yosuke Tanigawa, Fatima Rodriguez, Russ B. Altman, Nasa Sinnott-Armstrong, Manuel A. Rivas

## Abstract

Sex differences have been shown in laboratory biomarkers; however, the extent to which this is due to genetics is unknown. In this study, we infer sex-specific genetic parameters (heritability and genetic correlation) across 33 quantitative biomarker traits in 181,064 females and 156,135 males from the UK Biobank study. We apply a Bayesian mixture model, Sex Effects Mixture Model, to Genome-wide Association Study summary statistics in order to (1) estimate the contributions of sex to the genetic variance of these biomarkers and (2) identify variants whose statistical association with these traits is sex-specific. We find that the genetics of most biomarker traits are shared between males and females, with the notable exception of testosterone, where we identify 119 female and 444 male-specific variants. These include protein-altering variants in steroid hormone production genes (*POR*, *CYP3A43*, *UGT2B7*). Using the sex-specific variants as genetic instruments for Mendelian Randomization, we find evidence for causal links between testosterone levels and height, body mass index, waist circumference, and type 2 diabetes. We also show that sex-specific polygenic risk score models for testosterone outperform a combined model. Overall, these results demonstrate that while sex has a limited role in the genetics of most biomarker traits, sex plays an important role in testosterone genetics.

## Introduction

Sex differences have been documented in many phenotypes and diseases (Ober, Loisel, and Gilad 2008). Across quantitative traits, men and women typically have overlapping distributions with different means, examples of these traits include height and body mass index (BMI) (Winkler et al. 2015). Previous studies have demonstrated that some of this difference is due to sex-specific genetic factors (Ober, Loisel, and Gilad 2008; Khramtsova, Davis, and Stranger 2019). Genome-wide Association Studies (GWAS) are increasingly used to identify variants that contribute to sex differences, and recently, gene-by-sex interactions have been identified across many phenotypes, including in anthropometric traits (Rask-Andersen et al. 2019; Randall et al. 2013), irritable bowel syndrome (Bonfiglio et al. 2018), and glioma (Ostrom et al. 2017).

Examination of blood and urine laboratory biomarker levels reveal sex differences (Sinnott-Armstrong et al., n.d.); however, it is unknown to what extent these differences are related to underlying differences in the genetic architecture in men and women versus environmental effects. Narrow-sense heritability (h^2^) is the fraction of phenotypic variability explained by additive genetic variance. Heritability analysis helps to estimate the extent to which a trait can be predicted based on genetic risk rather than environmental factors. Initially, heritability was estimated from family studies using linkage; but now, with the increasing availability of genome-wide data, heritability is often estimated using genetic variants such as single nucleotide polymorphisms (SNP). SNP-based heritability represents the fraction of phenotypic variance due to additive effects from common genetic variation (Gamazon and Park, n.d.; Yang et al. 2017). Methods for estimating SNP-based heritability include LD-score regression, GREML (genomic related matrix restricted maximum likelihood), Haseman-Elston Regression, and the moment-matching approach (Bulik-Sullivan, n.d.; Hill 1978; Ni et al. 2018; Speed and Balding 2019). These methods are applied to a sample of unrelated individuals in order to quantify the proportion of phenotypic variance explained by all genetic variants in the GWAS.

At a trait-level, we can use the sex-specific heritabilities and the genetic correlation between males and females to examine what fraction of the genetics of that trait is shared between sexes. The UK Biobank is a prospective population-based study of 500,000 individuals that includes both genetic and phenotypic data, allowing for rich SNP-based estimation of heritability (Sudlow et al. 2015). While most traits do not show sex effects on heritability(Ge et al. 2017),^(Stringer, Polderman, and Posthuma 2018)^, previous studies have documented these differences are found in a subset of traits; for example, one study used traditional methods to identify differences in 11 out of 20 quantitative traits in the Hutterite population (Pan, Ober, and Abney 2007), and two recent studies found differences in fat distribution (Pulit et al. 2019) and anthropometric traits in the UK Biobank population with SNP-based methods (Rawlik, Canela-Xandri, and Tenesa 2016). These results demonstrate that sex differences in genetic effects exist, but to date, no analysis of these sex differences across biomarkers has been conducted.

Here we present an approach for estimating the extent to which genetic effects are correlated between sexes and identifying the proportion of relevant variants that have shared effects versus effects that are specific to each sex. We apply this approach to blood and urine biomarker data from the UK Biobank to examine sex differences in genetic effects, and find differences primarily in the genetic determinants of testosterone level. Furthermore, we use these identified sex differences to provide hypotheses about biological mechanisms including (1) examination of protein-altering variants and tissues where these genes are selectively expresssed, (2) causal inference using Mendelian Randomization to assess relationships between testosterone and other traits, and (3) improved genetic risk prediction models for testosterone.

## Results

### Sex Effect Mixture Models

We built a two-component Bayesian Sex Effect Mixture Model (SEMM) for estimating the contributions of sex to genetic variance using GWAS summary statistics (**Figure 1**). The model contains a null component, for variants that do not contribute to the trait, and a non-null component, for variants related to the trait, representing the genetic contribution to that trait. Variants driving male and female traits in the non-null component are modeled as two-dimensional vectors drawn from a multivariate normal distribution with a variance-covariance matrix that can be used to estimate the genetic correlation between sexes. To assess whether our approach obtains reliable statistical summaries on real GWAS data we applied the two-component SEMM to traits studied in (Rawlik, Canela-Xandri, and Tenesa 2016) and obtained comparable heritability and genetic correlation estimates (**Supplemental Figure 1**, **Supplemental Table 1**).

**Figure 1.**
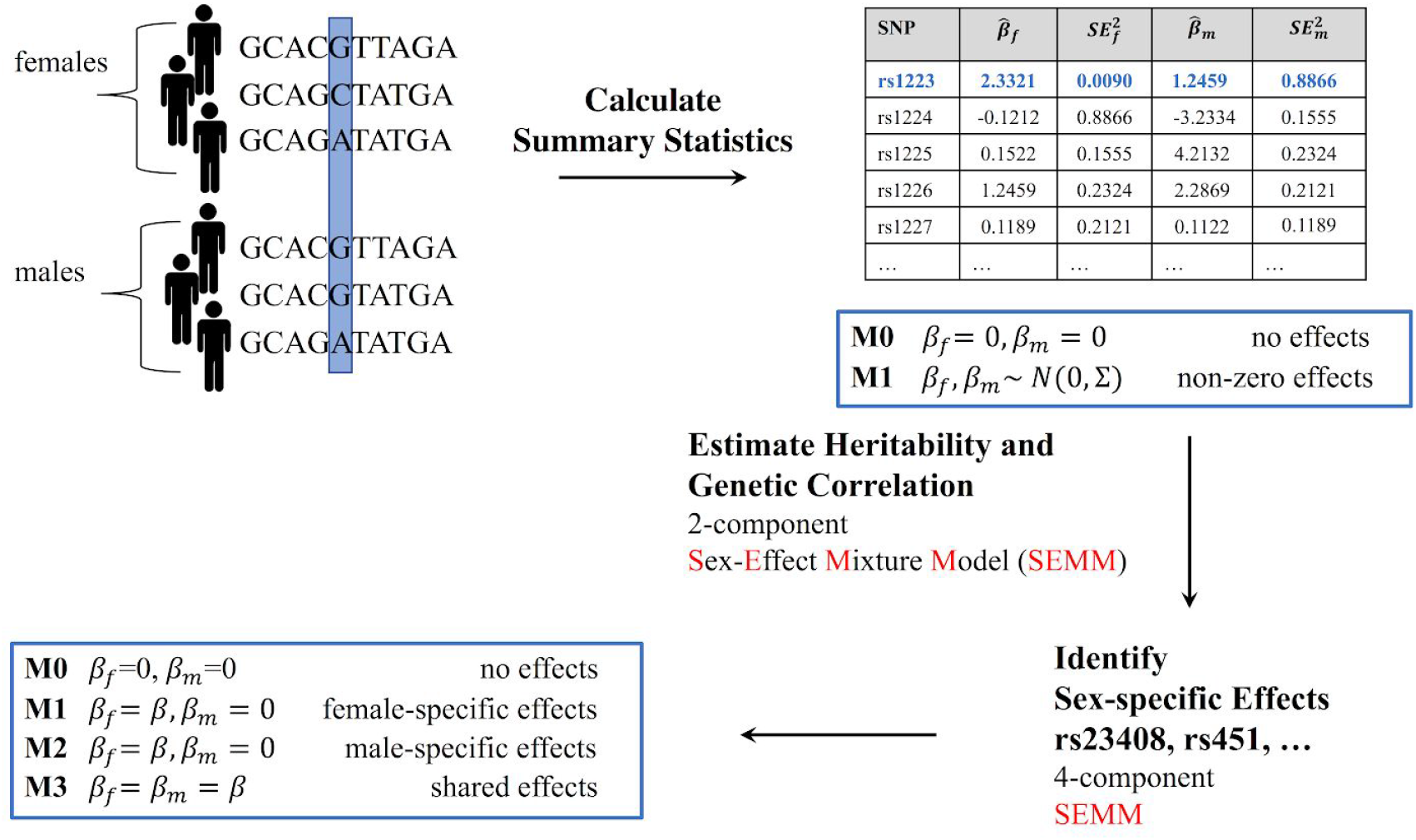
Schematic overview of Sex-Effect Mixture Model. We prepared a dataset of 33 serum and urine biomarkers from 358,072 individuals in UK Biobank. We calculated GWAS summary statistics from males and females separately, so that for every trait, we had an effect estimate (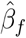 and 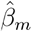) and standard error for each variant in each sex. We use a two-component Bayesian Sex-Effect Mixture Model (SEMM), with no effect and non-zero effect components, to estimate SNP-based heritability and genetic correlation between males and females for each biomarker. A four-component extension of the SEMM contains two additional components for separate male and female effects. This model allows us to distinguish between four cases: genetic variants that have no effect (illustrated as M0), genetic variants that have a stronger association with the trait in females or males (M1 and M2), and genetic variants that have similar effects in females and males (M3).

We extended our two-component SEMM to a four-component model to identify genetic variants with different effects in males and females (**Figure 1**). To do so, we add two components; one for detecting genetic variants that have stronger effects in males and the other for detecting genetic variant with stronger effects in females. Similar to the two-component model, the four-component SEMM also contains a no-effect and shared-effect component. Through fitting this model, we are able to separate genetic variants with different effects between men and women from those with shared effects. A “shared” effect would mean that the same genetic variant or set of variants have the same effect in males and females; “sex-specific” effects would indicate that the effect of these variants in males is different from that in females -- e.g. this variant is associated with higher lab test values in females but not in males.

To demonstrate the efficacy of the four-component SEMM on real data, we applied the model to four traits, waist-hip ratio, arm-fat-ratio, leg-fat-ratio, and trunk-fat-ratio with previously identified sex-specific genetic effects (Rask-Andersen et al. 2019)(Randall et al. 2013). We identified 371, 528, 840, and 1,221 genetic variants that had significantly stronger associations in females in waist-hip-ratio, arm-fat-ratio, leg-fat-ratio, and trunk-fat-ratio respectively. In males, only 13 variants were found in arm-fat-ratio (estimated False Discovery Rate (FDR) 4.7-6.6% across all traits and sexes, see **Supplemental Tables 2a-c** for numbers of variants, the false discovery rates of these estimates, and the estimated model parameters). Included in the female-specific waist-hip ratio variants were genetic variants proximal to four of the six previously reported genes (*COBLL1*/*GRB14*, *VEGFA*, *PPARG*, *HSD17B4*). Fat ratio variants were proximal to one of the male- and fifty-four of the female-specific genes previously identified, indicating that we capture known sex-specific signal. (see **Supplemental Table 2d** for the overlap with (Rask-Andersen et al. 2019) and **Supplemental Tables 3a-c** for the full lists of variants).

### Sex-differential heritability

We applied the two-component SEMM to 33 biomarkers in the UK Biobank previously described in (Sinnott-Armstrong et al., n.d.) in order to estimate the male and female heritabilities and the shared genetic effects between sexes for each trait (**Methods** and **Supplemental Table 4** for a full list of these traits. While a large fraction of biomarkers had overlapping heritability estimates, we found sex differences in the heritability of 17 of 33 biomarkers, including testosterone, IGF-1, Non-albumin protein, SHBG, total protein (higher in males than females), Apoplipoprotein B, C-reactive protein, cholesterol, creatinine, Cystatin C, eGFR, Gamma glutamyltransferase, HDL-C, LDL-C, Potassium in urine, Sodium in urine, urate (higher in females than males, **Figure 2A**). Of these, Cholesterol, Creatinine and Sodium in Urine, LDL, Testosterone, and Urate showed greater than 1.3 fold differences. For the majority of traits, the between-sex genetic correlations were close to 1.0, which indicates shared additive genetic effects between males and females (**Figure 2B**). By contrast, for testosterone, we estimated a genetic correlation of only 0.120 (95% HPD 0.0805 to 0.163), indicating that the genetic effects in males and females are largely non-overlapping (see **Supplemental Table 5** for estimated heritabilities and genetic correlations for all biomarker traits).

**Figure 2.**
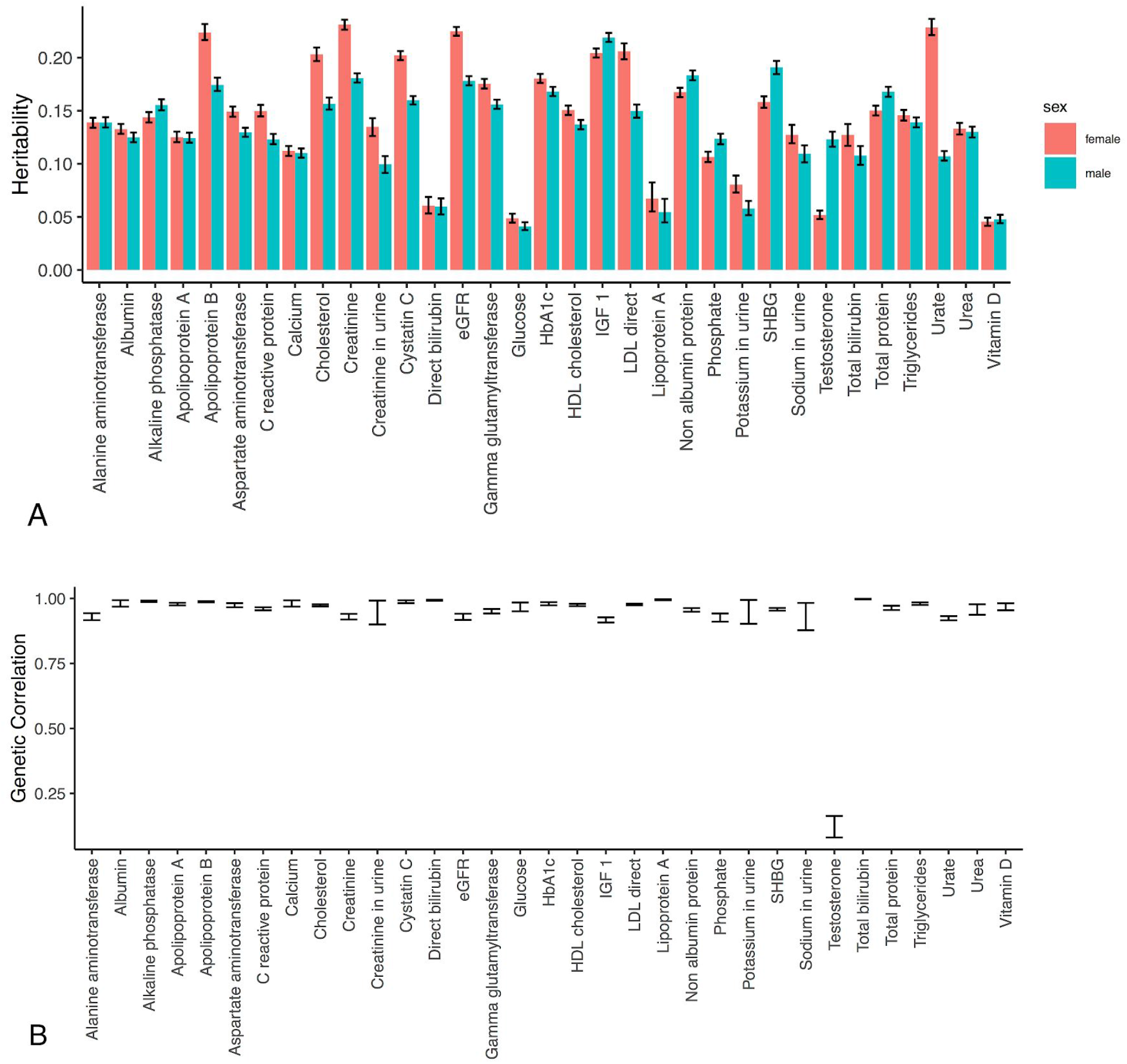
Heritability and genetic correlation of biomarkers between females and males. **A)** SNP-based heritability estimates of 33 biomarkers for females (red) and males (cyan). **B)** Correlation of genetic effects between males and females for 33 biomarkers. Error bars in both panels indicate the 95% highest posterior density.

The heritability of a particular trait can vary across the lifetime, as genetics may explain more or less of the variation in that particular trait. Previous studies have found that pre- and post-menopausal women have different heritability for BMI, waist and hip measures (Kelemen et al. 2010), and lipid biomarkers (Middelberg et al. 2002). To examine this across biomarkers in the UK Biobank population, we applied our two-component SEMM to GWAS summary statistics for pre- and post-menopausal women. We found that the genetic correlations between pre- and post-menopausal women were generally close to 1.0, and all traits had higher or equivalent within-sex genetic correlations (between pre- and post-menopausal women) when compared to between-sex correlations of either group with men (**Figure 3**, **Supplemental Table 6**).

**Figure 3.**
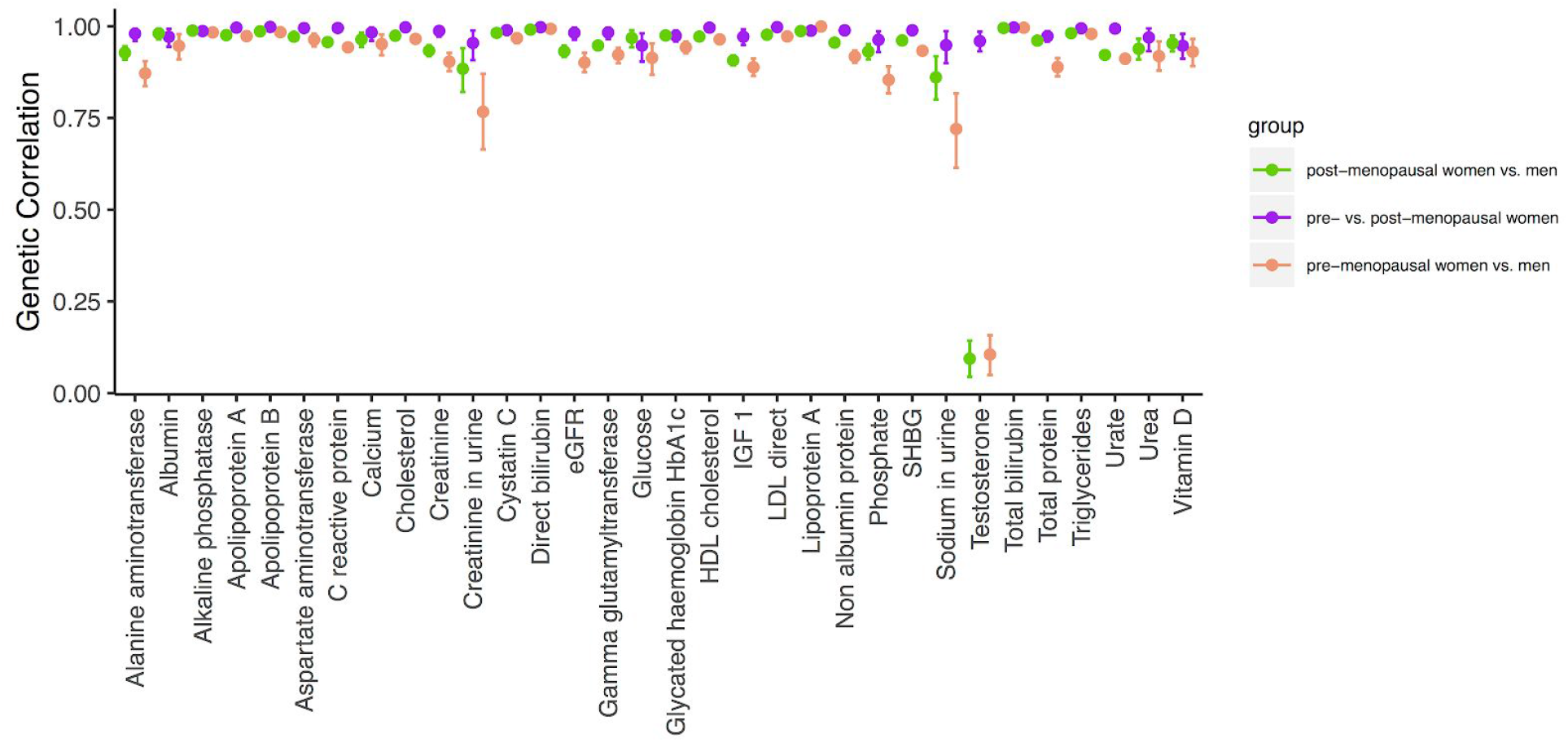
Genetic correlation and menopausal status. We examined whether the genetic correlation was affected by menopausal status. The genetic correlation within women (pre- vs. post-menopausal, in purple) was higher or equal to than either that between both post-menopausal women and men (green) and pre-menopausal women and men (orange). Error bars indicate the 95% highest posterior density.

### Identification of genetic variants with sex-specific effects

We applied our four-component SEMM to all 33 biomarkers to identify genetic variants with sex-specific and shared effects. In total, our analysis found 26,487 variants with effects on the traits of interest (**Methods**, **Supplemental Table 7**). As expected, the majority (25,880) of these variants showed shared effects between sexes, and most traits had few or no sex-specific variants. We identified 146 and 463 genetic variants with sex-specific effects in females and males respectively, the bulk of them corresponding to sex-specific genetic effects on testosterone levels (81.5% and 95.9% respectively; see **Figure 4** to visualize the effect sizes of these variants and **Supplemental Table 8** for a full list of genetic variants).

**Figure 4.**
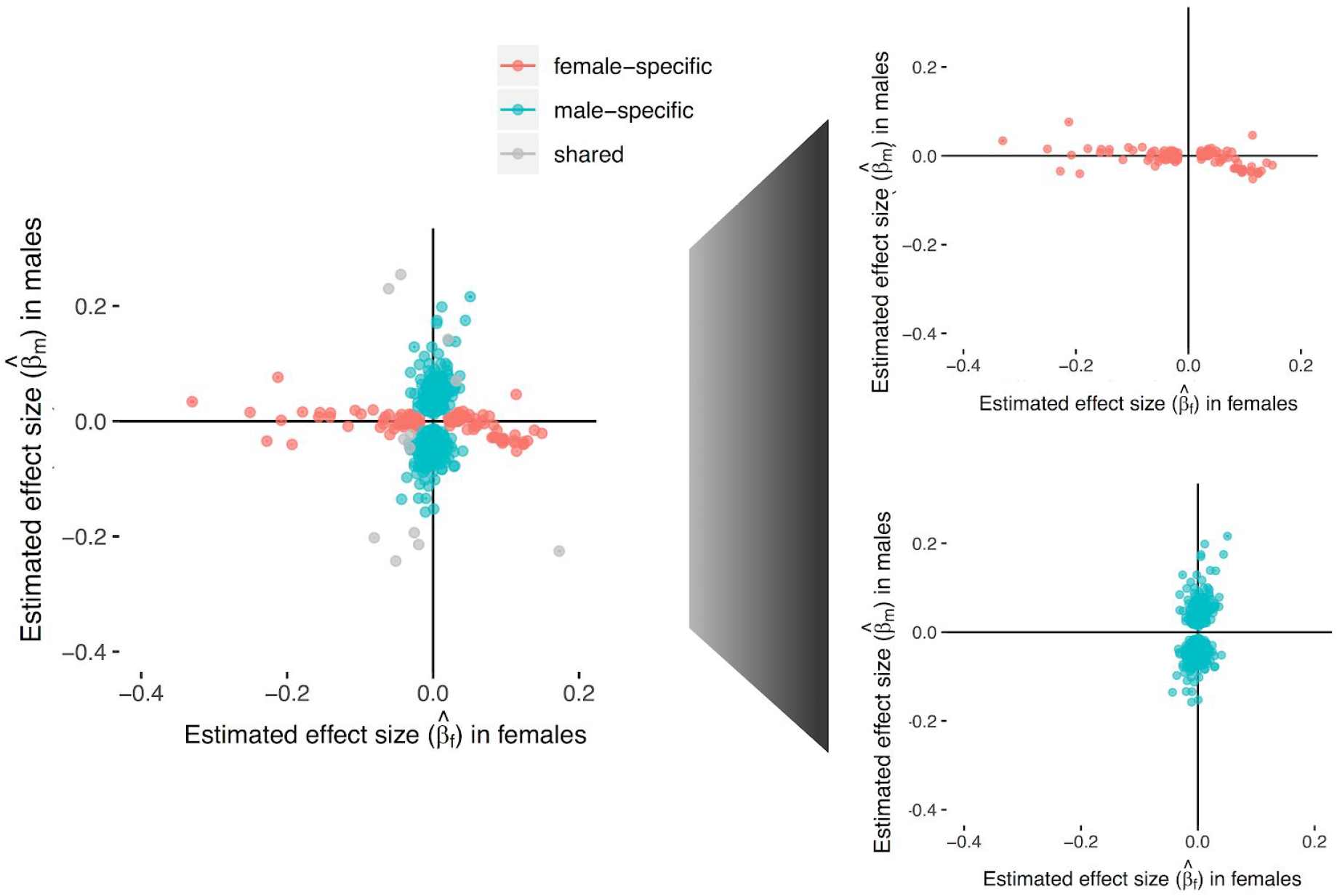
Identification of genetic variants with sex-specific effects on testosterone levels. Estimated effect sizes for genetic variants with non-zero effects on testosterone are shown (x-axis, estimated effect size in females; y-axis, estimated effect size in males). Blue dots correspond to variants that belong to the “Male-specific” effect component; red corresponds to the “Female-specific” effect component; and gray dots correspond to genetic variants that belong to the “Shared” effect component of SEMM.

Of the variants associated with testosterone, 55 male-specific variants, 1 female-specific, and 1 shared variant are located on the X chromosome, this indicates enrichment of X chromosomal variants in male testosterone genetics and is consistent with previous reports (Ohlsson et al. 2011) In addition, we recover variants in known testosterone-related genes (*AR*, *JMJ1DC*, and *FAM9B*) specifically in males, but not in females.

Previous studies of testosterone genetics have focused on males; here we identify multiple female-specific variants with strong positive or negative effects on testosterone. We examined the subset of these variants that encode missense variants in protein-coding regions; this included 17 variants in females and 59 variants in males (listed in **Supplemental Table 8a** and **8b** respectively). In females, these variants include *rs149048452* in *STAG3* (*β*=−0.33 and p=8.94×10^−9^ in females) and rs17853284 in *POR* (*β*=−0.23 and p=8.91×10^−15^). *STAG3* encodes a part of the cohesin complex involved in meiosis, previously researchers have found that mutations in this gene are associated with premature ovarian failure (Le Quesne Stabej et al. 2016). *POR* encodes a cytochrome p450 oxidoreductase; deficiencies in this enzyme have been associated with amenorrhea, disordered steroidogenesis, and congenital adrenal hyperplasia (Idkowiak et al. 2005). Many female-specific missense variants are located in genes associated with steroid hormone production (*LIPE*, *POR*, *CYP3A43*, *UGT2B7*) or gamete formation (*STAG3*, *MCM9*, *TSBP1*, *ZAN*); although *ZAN* and *TSBP1* encode the sperm zonadhesin protein and testis-expressed protein 1 respectively. To our knowledge, these associations with testosterone levels have not been discovered previously, and may help us to understand testosterone levels in women.

To assess the replicability of these findings, we also examined the effect sizes of sex-specific variants in a held-out cohort of Non-British White individuals (n=10,546 males and 14,269 females) (**Supplemental Figure 3**, **Supplemental Table 9**). For testosterone, we see a pattern of high concordance between the discovery and validation effect size estimates for sex the variants are associated with (R^2^ > 0.56) but not for the opposite sex (R^2^ < 0.05); this is also seen in the anthropometric traits we examined. The results indicate that variants we identify with SEMM have consistent but attenuated effects in a held-out cohort, which is expected due to winner’s curse (Palmer and Pe’er 2017).

#### Sex-specific genetic effects on testosterone are selectively expressed in liver

Given the substantial number of sex-specific variants associated with testosterone levels, we sought to understand the impact of those variants on the tissue-specific expression of the nearby genes. Tissue-Specific Enrichment Analysis (TSEA) leverages gene expression data from the Genotype-Tissue Expression (GTEx) dataset to calculate the enrichment of tissue genes within a provided gene list. We applied TSEA to the genes containing variants associated with testosterone levels identified by SEMM. In the male component genes, we found enrichment for liver-specific genes (Benjamini-Hochberg corrected p = 3.91×10^−7^ respectively, **Supplemental Figure 4**). By contrast, the female component genes showed no significant enrichment after correction for multiple hypothesis testing (Benjamini-Hochberg corrected p > 0.1). In men, low serum testosterone levels are associated with cirrhosis and non-alcoholic fatty liver disease (NAFLD), with decreasing levels corresponding to increased disease severity (Sinclair et al. 2015; Mody et al. 2015; Yim et al. 2018). Previous studies also find this inverse relationship in post-menopausal women (Sinclair et al. 2015; Mody et al. 2015; Yim et al. 2018); however, women with polycystic ovarian syndrome, which is associated with increased androgens, have increased risk of NAFLD independent of obesity status (Kim et al. 2017).

### Mendelian randomization of sex-specific genetic effects

After identifying genetic variants with sex-specific effects on testosterone levels, we used Mendelian Randomization (MR) to examine whether these biomarkers are causally related to disease outcomes or other commonly measured traits. The intuition is that if a genetic variant is associated with differing levels of the biomarker, this provides a natural experiment, and we can examine whether the predicted variance in that biomarker based on the genetic variant is associated with the outcome variance, which indicates a causal effect. Recently, an MR study found evidence for a causal link between higher testosterone levels and cardiovascular disease in men using *JMJD1C* variants in the UK Biobank population (Luo et al. 2019) (Twelve of the genetic variants assigned to our male-specific component are located in *JMJD1C*). Additional studies in men used testosterone variants as instruments to test for a causal relationship of testosterone on cognition (Zhao et al. 2016) and BMI (Eriksson et al. 2017), but did not find evidence of associations. While there is an extensive focus on testosterone levels in men, (Luo et al. 2019) and (Schooling et al. 2018) performed the only testosterone MR study that included women. The authors found evidence of causal links between testosterone and a variety of cardiovascular risk factors in men or men and women combined, but not in women. However, this analysis was limited by their genetic instruments; a previous GWAS in post-menopausal women did not find genetic variants associated with testosterone levels (Prescott et al. 2012).

The sex-specific testosterone variants identified by our analysis provide a unique opportunity to further examine the causal effects of testosterone levels in both men and women. We aggregated a total of 10 outcomes (**Supplemental Table 10**), including anthropometric traits (height, BMI, waist circumference [WC], and hip circumference), disease outcomes (heart disease, stroke, and diabetes), and sex-specific traits (ages at menarche and menopause, prostate cancer, **Methods**), and assessed the causal effects of these variants using the Inverse-Variance Weighted (IVW) Method (van der Plaat et al. 2019; Bowden et al. 2017) (see **Figure 5** **and Supplemental Table 11** for the results).

**Figure 5.**
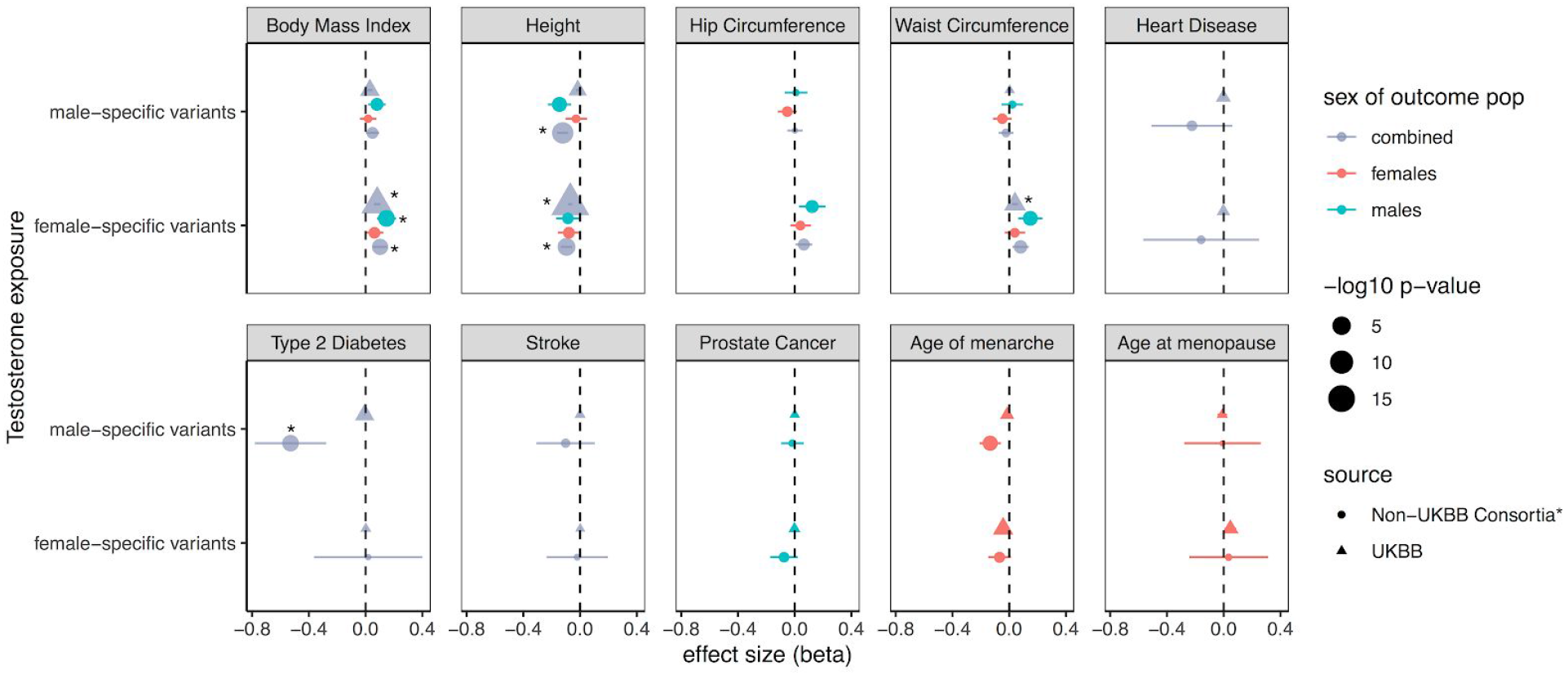
Results of Mendelian Randomization tests with sex-specific testosterone variants as instruments on all analyzed outcomes. This plot shows the Mendelian Randomization results for all traits, each trait is shown separately. Effect sizes (betas) are estimated using either the female or male-specific variants as instrumental variables for testosterone exposure. 95% confidence intervals are shown for each estimate. Points are colored by the sex of the outcome population (blue for males, red for females, and gray for combined), with size indicating the −log10 p-value, and shape showing the source of the GWAS statistics for the trait (triangles for UKBB = UK Biobank, circles for all others). *The list of Non-UK Biobank consortia traits are shown in **Methods** and **Supplemental Table 10**.

We found that testosterone levels showed evidence of a causal association with BMI and WC using female-specific variants as instruments with estimated effects consistent with higher testosterone increasing BMI and WC (p=8.0×10^−5^, 8.6×10^−5^; β = 0.10, 0.04; standard error = 0.026, 0.01 respectively **Supplemental Figure 5**). A previous MR study on the UK Biobank examined testosterone for causal effects on waist circumference and BMI, but did not find evidence of an association (Schooling et al. 2018); however, it is possible we are able to find these associations because we used sex-specific genetic instruments. Both female and male variants showed evidence of a causal association with height (p=3.7×10^−6^ and 7×10^−10^), with higher testosterone associated with decreased height (β =−0.96 and −0.12, SE=0.021, 0.020). This is in contrast to previous findings of a positive relationship between height and testosterone levels at a population level (Schooling et al. 2018; Handelsman et al. 2015) and in a previous MR study (Schooling et al. 2018; Handelsman et al. 2015). For all of these associations, we observe similar effects in both the UK Biobank and GIANT datasets, and using MR Egger and IVW (**Supplemental Table 11** contains a full list of the effect sizes and p-values for all MR tests).

Male-specific testosterone levels show evidence of an association with type 2 diabetes (p=3.6×10^−5^); higher testosterone is related to type 2 diabetes risk reduction (β =−0.52, SE=0.12) using data from the combined DIAGRAM and MetaboChip study (Morris et al. 2012). This association was found using the IVW method; MR Egger estimates indicate the relationship is in the reverse direction and is not significant (*β*=0.81, SE=0.59, p=0.18). Further work is required to examine if this discrepancy is due to confounding. Several longitudinal studies have shown that low levels of testosterone predict the later development of type 2 diabetes or metabolic syndrome (Stellato et al. 2000; Antonio et al. 2015). However, in a small clinical trial of 88 individuals the effect of testosterone treatment on glucose metabolism was assessed finding no evidence of an effect (Gianatti et al. 2014). This study contradicted earlier data from the TIMES2 clinical trial study (n=220) showing evidence of testosterone replacement therapy effects on insulin resistance and symptoms of hypogonadal men with type 2 diabetes or metabolic syndrome.

### Sex-specific multivariate polygenic risk prediction

Motivated by the sex differences in testosterone genetics, we tested whether constructing sex-specifc polygenic risk scores (PRS) would have better performance for predicting testosterone levels than a sex-combined model. We applied batch screening iterative lasso
(BASIL) to train multivariate penalized regression models for males and females independently (“sex-specific” model) and both males and females combined (“combined” model, **Methods**) (Qian et al. 2019). In male and female individuals separately, we compared the sex-specific to the combined models to see if building a separate model for each sex affected our ability to predict covariate-adjusted testosterone levels. We evaluated the model on a held-out test set, and found that the sex-specific PRS and the combined PRS are consistent (`ρ` = 0.59 and 0.60, both with p < 2.2×10^−16^ for the correlation between the combined model and each of male and female models, respectively, **Supplemental Figure 6**). Still, we found the sex-specific PRS models have improved predictions of covariate-adjusted testosterone in the sexes they were trained on (R^2^ = 0.31 and 0.18 for male and female, respectively) over the combined model (R^2^ = 0.21 and 0.13 for male and female, respectively). Sex-specific PRS trained for the opposite sex have low predictive performance (R^2^ = 0.020 and 0.023 for female-specific PRS evaluated on males and male-specific PRS evaluated on females, respectively). Overall, these results highlight the benefits of sex-specific polygenic prediction for testosterone levels (**Figure 6**).

**Figure 6.**
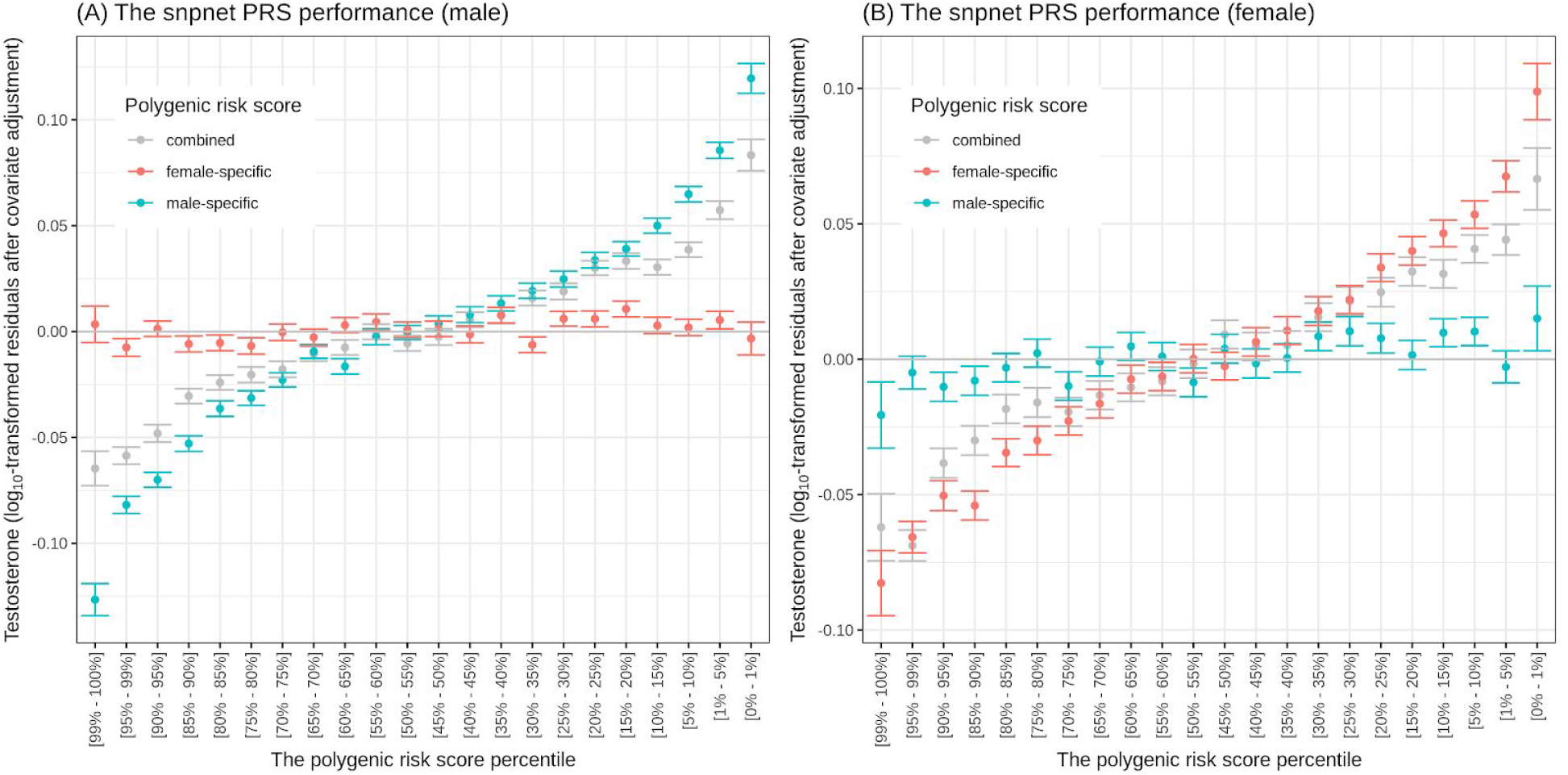
Sex-specific polygenic prediction. The mean testosterone values are shown (y-axis) across stratified risk bins (x-axis) based on sex-specific PRS (blue for male PRS and red for female PRS) or the combined PRS (gray) for
male and (B) female individuals. Mean values and PRS scores are calculated on a held-out test data set of unrelated White British individuals and the testosterone are shown as log_10_-transformed values of residuals after the covariate adjustment (**Methods**). The error bars represent standard errors. The PRSs are computed with 8236, 7169, and 7320 genetic variants for male-specific, female-specific, and the combined model, respectively.

## Discussion

While known sex differences in biomarker levels have been reported, the extent to which these differences are encoded in the genome is not known. To answer this question, we studied the genetics of 33 biomarkers in UK Biobank males and females using a two and four-component Bayesian Mixture Model, SEMM. SEMM has the benefit of both identifying the underlying genetic architecture and identifying genetic variants with shared and sex-specific effects. For the majority of the traits we analyzed we do not see strong sex differences in genetic effects, which is expected and been previously documented in the literature (Stringer, Polderman, and Posthuma 2018). Namely, the genetics of these traits are shared (as indicated by genetic correlations close to one), the traits have similar heritability, and most of these traits have no variants with sex-specific effects.

By contrast, we found little overlap between males and females in the genetics of testosterone levels. In addition to finding significant sex differences in genetic architecture, we also identified over five hundred genetic variants with male- or female-specific effects on testosterone levels. Because of the male-female differences in testosterone genetics, we examined the subset of protein-altering variants and the tissue-specific expression patterns of genes with variants that have sex-specific genetic effects. The protein-altering variants associated with female-specific effects testosterone include variants in genes associated with steroid hormone production and gamete production. Tissue-specific enrichment analysis reveals that the genes proximal to these sex-specific variants are enriched in liver in males but not females. It is hypothesized that this relationship between testosterone and liver disease may have different etiology in men and women, and may also be mediated by interactions between testosterone, insulin resistance, obesity, and metabolic syndrome (Yim et al. 2018). Additionally, we built sex-specific polygenic risk models, which showed improved predictive performance over a sex-combined model.

We used Mendelian Randomization to assess whether testosterone may be causally implicated in a broad range of diseases and phenotype measurements. In this study, we found evidence of MR associations of testosterone with both BMI and waist circumference using female-specific variants associated with testosterone levels as instruments, height using both male and female variants, and type 2 diabetes using male-specific variants. The associations of testosterone with BMI and waist circumference indicate an increasing effect of testosterone on both traits, which is consistent with the hypothesis that testosterone is involved in obesity and metabolic syndrome. However, these relationships have not been previously reported from MR; it is possible that we are able to identify these associations because the female sex-specific genetic variants represent a different set of genetic instruments than have been previously used. The evidence of an association with height shows a decreasing effect of testosterone in height, which is surprising, as previous studies have indicated that higher testosterone may be associated with higher stature (Schooling et al. 2018; Handelsman et al. 2015). However, testosterone is sometimes used as a therapy for tall males with delayed puberty, and results in accelerated initial growth but overall stunting of stature (Zachmann et al. 1976). Further work is required to understand this association. Finally, the potential causal relationship with type 2 diabetes supports the hypothesis that testosterone treatment for reducing diabetes risk in men may be a worthwhile approach; however, this must be taken with caution because the relationship was seen using the IVW but not MR Egger method. A current study, T4DM, is underway to examine whether testosterone treatment can prevent type 2 diabetes in men who have prediabetes and low testosterone (Bracken et al. 2019).

SEMM is made publicly available as an R package. Additionally, the inference results are available for visualization as a web application in the Global Biobank Engine (McInnes et al. 2019), and all the sex-specific genetic variants are included in the Supplemental tables (**Supplemental Tables 3** and **8**). While we applied this method to examine sex differences in genetic effects, SEMM could also be used to for any other type of binary gene-covariate analysis.

Our understanding of male and female reproductive health is limited and evolving (Yamin and Boulanger 2014). We anticipate that further aggregation of UK Biobank data in these areas combined with the analysis of the genetic architecture of readily available traits across sexes will help identify the extent to which genetic effects that impact health outcomes are shared and sex-specific. Testosterone is frequently thought of as a male sex hormone because of its higher levels in men and involvement in the development of the male reproductive tract and secondary sex characteristics. However, females also produce testosterone, albeit at lower levels. Elevated testosterone levels in women have been associated with polycystic ovarian syndrome, insulin resistance, dyslipidemia, and hypertension (Amer 2009; Haring et al. 2011; Mody et al. 2015).

Previous work examining the genetics of testosterone in females did not find associations (Prescott et al. 2012) and previous Mendelian Randomization studies have been limited by the lack of known testosterone variants in women (Schooling et al. 2018). Our analysis expands on and addresses this issue by using a larger population, carefully adjusted biomarkers, and our SEMM method to identify these variants. Our results demonstrate that the genetics of testosterone levels is complex and highly polygenic in both males and females. Further, our work highlights the importance of also examining female variability in testosterone levels, and of considering sex as a variable in these analyses. We anticipate that future analyses that include rare genetic effects, more diverse population cohorts, and improved integration with reproductive health outcomes will improve our understanding of the translational impact of these findings.

## Author Contributions

M.A.R. conceived and designed the study. E.F. and M.A.R. designed and carried out the statistical and computational analyses. Y.T. performed PRS analysis. E.F., Y.T., N.S.A, and
M.A.R. carried out quality control of the data. The manuscript was written by E.F. and M.A.R. All authors provided feedback on the manuscript. M.A.R. supervised all aspects of the study.

## Acknowledgments

This research has been conducted using the UK Biobank resource. We thank all the participants in the UK Biobank study. E.F. is supported by the NIH NLM F31 Fellowship F31LM013053. R.B.A is supported by the NIH GM 102365, LM 005652 and the Chan Zuckerberg Biohub. Y.T. is supported by a Funai Overseas Scholarship from the Funai Foundation for Information Technology and School of Medicine at Stanford University F.R. is supported by a career development award from the National Heart, Lung, and Blood Institute, National Institutes of Health (1K01HL144607). The primary and processed data used to generate the analyses presented here are available in the UK Biobank access management system (https://amsportal.ukbiobank.ac.uk/) for application 24983, “Generating effective therapeutic hypotheses from genomic and hospital linkage data” (http://www.ukbiobank.ac.uk/wp-content/uploads/2017/06/24983-Dr-Manuel-Rivas.pdf), and the results are displayed in the Global Biobank Engine (https://biobankengine.stanford.edu). We would like to thank Stanford University and the Stanford Research Computing Center for providing computational resources and support that contributed to these research results.

## Methods

### Genotype and phenotype data

We used genotype data from the UK Biobank dataset release version 2 and the hg19 human genome reference for all analyses in the study (Bycroft et al. 2018). To minimize the variability due to population structure in our dataset, we restricted our analyses to unrelated White British individuals based on the following four criteria reported by the UK Biobank in the file “ukb_sqc_v2.txt”:

1. used to compute principal components (“used_in_pca_calculation” column)
2. not marked as outliers for heterozygosity and missing rates (“het_missing_outliers” column)
3. do not show putative sex chromosome aneuploidy (“putative_sex_chromo-some_aneuploidy” column)
4. have at most 10 putative third-degree relatives (“excess_relatives” column).
5. White British ancestry (“in_white_British_ancestry_subset” column)

We subsequently focused on a subset of individuals with non-missing values for covariates and biomarkers as described below.

For validation, we used unrelated individuals from the Non-British White population, using a combination of genotype PCs provided by UK Biobank (−20 ≤ PC1 ≤ 40 and −25 ≤ PC2 ≤ 10) and their self-reported ethnicity in response to UK Biobank Field ID 21000.

We annotated variants using the VEP LOFTEE plugin (https://github.com/konradjk/loftee) and variant quality control by comparing allele frequencies in the UK Biobank and gnomAD (gnomad.exomes.r2.0.1.sites.vcf.gz) as previously described (DeBoever et al. 2018). We focused on variants outside of major histocompatibility complex (MHC) region (chr6:25477797-36448354) and performed LD-pruning using PLINK with “--independent 50 5 2” as previously described (DeBoever et al. 2018; Tanigawa et al., n.d.). We filtered for variants with Hardy-Weinberg Equilibrium < 10^−7^ and less than 1% missingness. For X chromosome variants, we used plink --xchr-model 2, which codes males as 0/2; this accounts for X inactivation in females, but many genes escape from inactivation.

For polygenic risk modeling, we additionally used copy number variation (CNV) data and HLA allelotype information from the UK Biobank (UK Biobank data field ID 22182) (Aguirre, Rivas, and Priest, n.d.; Bycroft et al. 2018). Using PLINK v1.90b6.7, we combined all 805,426 genotyped variants on arrays, 275,180 CNVs, and 362 allelotypes into a combined dataset consists of in total of 1,080,968 variants.

#### Anthropometric Traits

To demonstrate the utility of our method, we applied SEMM to anthropometric traits previously examined in (Rawlik, Canela-Xandri, and Tenesa 2016; Rask-Andersen et al. 2019) (UK Biobank field IDs for these traits are provided in **Supplemental Table 1**). Four traits were derived: arm-fat-ratio, leg-fat-ratio, trunk-fat-ratio, and waist-hip-ratio (WHR). WHR was calculated by taking the ratio of waist circumference (ID:48) to hip circumference (ID:49). Fat ratio traits were calculated as described in (Rask-Andersen et al. 2019) using impedance measures. Briefly, we took the fat mass for each body area: trunk (ID:23128), arm (ID:23124, 23120), or leg (ID:23116, 23112) -- for arm and leg we summed right and left together -- and divided by the total fat mass (ID:23100). For each of the anthropometric traits, we removed individuals missing data or with values outside six standard deviations of the mean.

#### Selection of Biomarker Traits

We focused on 33 biomarkers of the 38 biomarkers described in (Sinnott-Armstrong et al., n.d.) (UKBB field ID column in **Supplemental Table 4**). This included blood biochemistry assays (28 assays total) and urinalysis (4 assays total) and two derived blood biomarkers (Non-albumin protein and eGFR) described in (Sinnott-Armstrong et al., n.d.). Cholesterol, Apolipoprotein B, and LDL were adjusted by statins, as described below. Two of the blood biochemistry assays (Oestradiol and Rheumatoid Factor) and one urine assay (Microalbumin in Urine) were excluded because they had a large fraction of levels below the reported range, as described in (Sinnott-Armstrong et al., n.d.).

#### Statin identification and LDL adjustment

Statin adjustment was performed as reported in (Sinnott-Armstrong et al., n.d.).

#### Testosterone-related medications

From our analysis of testosterone levels, we removed individuals taking the following testosterone-related drugs: methyltestosterone (1140857656), finasteride (1140868550), dutasteride (1141192000), testosterone (1140868532), mesterolone (1140868526), and cyproterone (1140876638;1140884634;1141192344).

#### Covariate correction

For biomarker traits, raw UK Biobank measurements for all reported individuals (excluding out of range and QC failed measurements) were filtered for White British individuals, separated by sex, and fit with linear regression against 123 covariates. These included demographics (age, age^2^), population structure (the top 40 principal components and indicators for each of the assessment centers in the UK Biobank), temporal variation (indicators for each month of participation, with the exception that all of 2006 and August through October of 2010 were assigned a single indicator), socioeconomic status indicators (Townsend deprivation indices and interactions with age), BMI, WHR, the genotyping array used, and technical confounders (blood draw time and its square and interactions with age; urine sample time and its square and interactions with age; sample dilution factor; fasting time, its square, and interactions with age; and interactions of blood draw and urine sample time with dilution factor). The residual from this regression was inverse normal transformed using the Blom transform and then used as the tested outcome.

#### Phenotype definition for menopause

We used a stringent definition for dividing individuals into pre- vs post-menopause menopause categories. We relied on self-reported outcomes stating whether they had reached menopause (field ID: 2724) and their age at menopause (ID:3581).

- Pre-menopause: stated they have not reached menopause and are less than 60 years old
- Post-menopause: >2 years post menopause and had menopause after age 40

We also excluded anyone who had missing information or stated they were not sure whether they had menopause, and individuals who may have gone through surgical menopause (stated having had an oophorectomy ID:2834 or hysterectomy in ID:3591 or 2724). From the White British population, this resulted in 35,999 pre-menopausal women and 91,462 post-menopausal women (53,573 women excluded).

#### Outcome traits in Mendelian Randomization analysis

For Mendelian Randomization analysis, we used 10 outcome traits from MRBase (Hemani et al. 2018); a full list of these traits with their MR Base IDs is provided in **Supplemental Table 10**. For each trait, we used both summary statistics from the UK Biobank and from an additional consortia. We constructed this setup to both maximize the sample size (in many cases, UK Biobank had the largest population for that trait) and also to reduce the effect of winner’s curse to give false-positive results in the UK Biobank population because the testosterone variants were identified in that group. We selected four anthropometric traits and six disease traits. All UK Biobank traits were from Elsworth et al. Non-UK Biobank Consortia are as follows: GIANT for Body Mass Index, Height, Hip Circumference, and Waist Circumference (Randall et al. 2013); CARDIoGRAMplusC4D for Heart Disease (Nikpay et al. 2015); DIAGRAMplusMetaboChip for Type 2 Diabetes (Morris et al. 2012); ISGC for stroke (Malik et al. 2018); and ReproGen for Age at Menarche and Age at Menopause (Day et al. 2017).

### Summary statistic generation

Genome-wide association summary statistics were generated separately for males and females using PLINK v2.00aLM (2 April 2019). Age, genotyping array used, and the first four principal components were included as covariates. Univariate association analyses for single variants were applied to the phenotypes independently. For the 33 biomarker traits, the raw phenotypes were adjusted for the 123 covariates, and the analyses were performed on the residuals. Following summary statistic generation, alleles were flipped so that the effect size was always reported with respect to the alternate allele. Variants with missing standard errors or standard errors > 0.2 in either sex were also removed.

### Two Component SEMM – Estimating variance-covariance matrix for sex divided data

We specify a two-component mixture model consisting of a point mass centered at zero and a multivariate normal distribution. The components are described below:

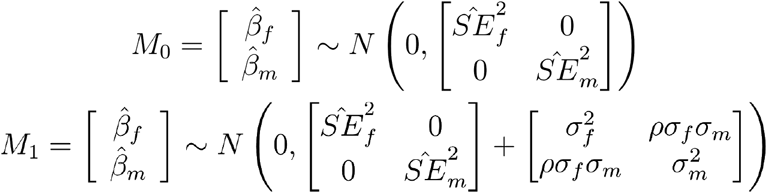

The model priors were set as follows:

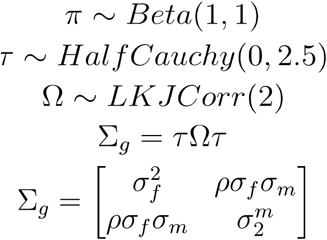

The likelihood across all variants is then formulated as:

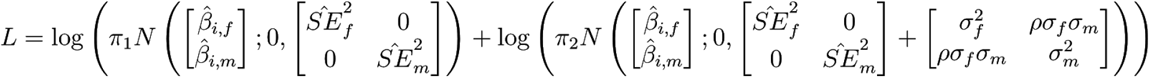

where *i* is the *i*th SNP.

We estimate π the proportion in each component, and Σ_*g*_, the genetic variance-covariance matrix for the non-null component. Ω provides the genetic correlation.

A Beta distribution centered at (1,1), was used for *π* in order to not favor assignment to either component. Priors were chosen for Σ_*g*_ based on suggestions from the STAN Manual (McElreath 2018). A half-Cauchy distribution with a small scale was chosen for τ to be a weakly informative scaling prior. The LKJ correlation distribution was chosen as a prior for Ω. This distribution is the identity correlation matrix (no correlation) at *v*=1, and as *v* increases for *v* > 1 the distribution shows less correlation between components and increasingly concentrates around the unit correlation matrix.

### Four Component SEMM - Identifying Sex-specific Genetic Variants

We formulated a mixture model with four components: (0) no effect, (1) female-specific effect, (2) male-specific effect, and (3) effects in both sexes.

This is described as follows, where *k* refers to the component. Here *β* ≠ 0:

*k* = 0, *β*_*m*_ = *β*_*f*_ = 0 No effect
*k* = 1, *β*_*m*_ = *β*, *β*_*m*_ = 0 Female-specific effect
*k* = 2, *β*_*f*_ = 0, *β*_*m*_ = *β* Male-specific effect
*k* = 3, *β*_*m*_ = *β*, *β*_*f*_ = *β* Effects in both sexes

For each component *k* = *i*, we have the following:

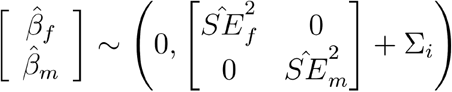

where:

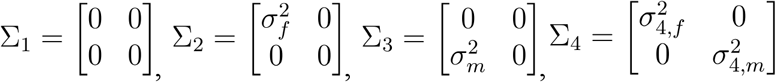

The priors are provided below:

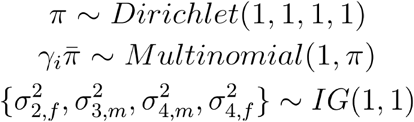

Here, we estimate π, the proportion in each component, and 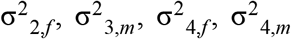, the non-zero variances for each component.

A Dirichlet prior centered at (1,1,1,1) was chosen for π in order to not favor any component at the start. InverseGamma(1,1) priors were used to cover a range of values expected for the nonzero variances σ^2^.

#### Assignment of Variants to Components

STAN does not assign samples to components in mixture models. In order to get this assignment, estimated parameters were used to assign variants to components. The probability of variant *i* in component *k* (i.e. γ_*i*_ = *k*) was modeled as a multinomial distribution with probability *p*_*i*_ where *p*_*i*,*k*_ is the probability of a variant with

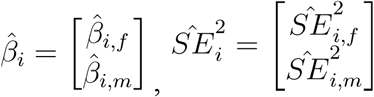

being in component *k* based on the model and estimated parameters^6^. This is formulated below:

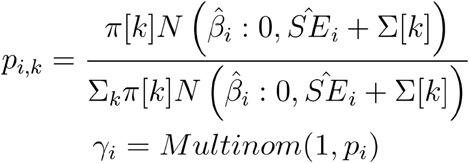

Variants were assigned to a non-null component if they had a posterior probability *p*_*i*,*k*_ > 0.8 of being in that component; otherwise, they were assigned to the null component.

#### False Discovery Rate (FDR) Calculation

To estimate the False Discovery Rate for the variants in a non-null component being associated with a trait, we first assigned the variants to components based on a posterior cutoff, as described above. For components with more than five assigned variants, we averaged the null posterior probabilities across all variants assigned to that component. We calculated the FDR for a range of posterior probability cutoffs (0.5-0.9) and used a cutoff of 0.8 in all subsequent analyses (see **Supplement Tables 2b and 7b** for the FDR estimates).

#### Computation of genetic correlation and heritability

Genetic correlations were estimated by Ω from the fit, the 95% highest posterior density interval (HPD) was given from the STAN fit of the parameter. Estimates for π and Σ are also extracted using the median value (50%) of the STAN fit for those parameters.

For each trait, the sex-specific heritability in males or females (*x*=*m* or *f*) was calculated by first assigning variants to components based on their posterior probability. Then using all variants assigned to the non-null component, heritability was estimated by the following equation, where *n*_1_ is the number of variants in the non-null component and π_1_ is the fraction in the non-null component.

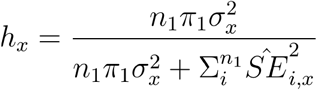

To create a 95% HPD interval, the posterior probability for each variant and the overall heritability was calculated as described above using the π and Σ estimates from each of the post-warmup draws. These estimates were then ordered, and the 2.5 and 97.5 percentile are reported as the 95% HPD interval.

#### Validation in Non-British White Cohort

We sought to assess the replicability of identifying sex-specific variants using the four-component model. In order to do so, we separately ran GWAS on only the sex-specific variants in the Non-British White population (n=10546 males and 14269 females). We focused on the anthropometric traits (WHR, leg-fat-ratio, arm-fat-ratio, trunk-fat-ratio) and testosterone because these traits have high numbers of sex-specific variants. We compared the variant effect size estimates in each sex between the discovery (British White) and validation (Non-British White) cohorts by calculating the R^2^ for using a linear model with the discovery as the independent variable and validation as the dependent variable for each set of sex-specific variants, using the male and female-specific effect estimates separately. Plots are shown for female- and male-specific variants in testosterone and female-specific variants in WHR in **Supplemental Figure 3a-c**. The R^2^ values for all five traits are included in **Supplemental Table 9**.

#### Computing Environment

Models were built and parameters estimated using STAN version 2.17.2 (Carpenter et al. 2017). Each STAN model was run with 4 chains, 200 warm-up iterations, and 800 total iterations. All models were checked for convergence as indicated by an Rhat value close to one. All other code for data processing and analysis was written in R (version 3.4.1).

#### Code Availability

The code to run SEMM is made publicly available as an R package on GitHub at https://github.com/rivas-lab/semm.

### Tissue-Specific Enrichment Analysis

We used the Tissue-Specific Expression Analysis tool to perform tissue enrichment analysis (the cell type version of this tool is described in (Xu et al. 2014). TSEA uses published RNA-seq data GTEx data (pilot data 2013-01-31), and calculates the enrichment of of a list of genes in the genes specific to each of twenty-five tissues. Briefly, the authors grouped data from 45 tissues (including sub-tissues) into 25 “whole-tissue” types by averaging the gene level read counts. They used their specificity index statistic (Dougherty et al. 2010), (Xu et al. 2014) to identify enriched transcripts for each of these tissues at three different thresholds (0.0001, 0.001, 0.01), where transcripts at the 0.0001 level are more specific to that tissue, and those at 0.01 are more general. A Fisher’s Exact test is then used to calculate a p-value for the enrichment of a set of genes in that set; this is Benjamini-Hochberg corrected to account for multiple tests. We used the genes proximal to sex-specific testosterone variants as input, and examined the tissue enrichment in males and females separately.

### Sex-specific multivariate polygenic prediction

To construct sex-stratified polygenic risk models using multivariate penalized regression, we applied batch screening iterative lasso (BASIL) implemented in the R snpnet package (Qian et al. 2019). We created a random split dataset of White British individuals in UK Biobank into 70% training (n = 236,005), 10% validation (n = 33,716), and 20% test (n = 67,430) sets, and used both training and validation sets for the analysis. The validation set was used to select the optimal lambda value that controls the sparsity. Focusing on individuals with non-missing phenotype values, we first took the log10-transformation of the testosterone value and defined covariate-adjusted phenotypes using sex-independent covariates (for male- and female-specific models) using a linear regression model. Similarly, a combination of the sex-independent covariates and an indicator variable “sex” (for combined model) was used to define the covariate-adjusted phenotype for the combined cohort of males and females after the log_10_-transformation. The list of covariates used in the polygenic prediction and their corresponding BETA for covariate correction is provided in **Supplemental Table 12** and covariate adjustment was performed independently for male-only, female-only, and the combined cohort. We used a combined genotype dataset consists of array-genotyped variants, copy number variants, and HLA allelotypes. Focusing on males (training n = 100,913; validation n = 14,594) and females (training n = 99,564; validation n = 14,049) in unrelated White British individuals, we independently trained sex-specific PRS models. As a baseline for comparison, we additionally trained the combined model using a cohort consists of both sexes (training n = 200,477; validation n = 28,643). We extracted non-zero regression coefficients (BETAs) from the selected optimal model and computed sex-specific polygenic risk scores using --score function implemented in plink version 2.00a2LM (26 Aug 2019).

To evaluate the performance of sex-specific PRS or the “combined” PRS, we stratified individuals in the test set (males n = 28,601; females n = 28,640; combined n = 57,241) based on the PRS bins and computed the mean value and standard error for each bin. The standard error was computed by dividing the sample standard deviation by the square root of the sample size in the bin.

To compare and evaluate the consistency of sex-specific PRS and the combined PRS, we wsimilarity using Spearman’s rank correlation (`ρ`) (**Supplemental Figure 6**). The PRSs in the plots are centered and scaled so that each has zero mean and unit variance.

### Mendelian Randomization

We used MR-Base to test for evidence for causal associations between testosterone and 10 outcomes of interest using the sets of female- and male-specific testosterone variants as instrumental variables (Hemani et al. 2018). We used clumping to prune variants for LD and performed the analysis with three methods: MR Egger, Inverse Variance Weighted, and Inverse Variance Weighted with fixed effects. For each of the outcomes, we used summary statistics from both a UK Biobank and non-UK Biobank source, we also used sex-divided outcomes for the four anthropometric traits for which they were available. The traits include: waist circumference (Shungin et al. 2015), hip circumference (Shungin et al. 2015), height (Wood et al. 2014; Randall et al. 2013), body mass index (Locke et al. 2015), age at menarche (Perry et al. 2014), age at menopause (Day et al. 2015), prostate cancer (Schumacher et al. 2018), heart disease (CARDIoGRAMplusC4D Consortium et al. 2013), type 2 diabetes (Morris et al. 2012), and stroke (Malik et al. 2016). See **Supplemental Table 10** for more information about these traits and their MR-Base ids. We used a Bonferroni correction to account for multiple tests (p-value threshold = 0.05/168=2.98×10^−4^).

