## Supplemental Figures and Information for "Sex-specific genetic effects across biomarkers"

#### Supplemental Tables

Supplemental Tables are included as excel files, captions are below.

**Supplemental Table 1. SEMM Genetic Parameter Estimates for Anthropometric traits.** For each trait (trait, UK Biobank data field ID [UKBB ID], parameter fits (*pi*[0] is the null and *pi*[1] is the non-null fraction, and *Sigma* is the variance-covariance matrix, see **Methods**) and derived quantities, including heritability (*hf* = female-specific heritability, *hm* = male-specific heritability) and between-sex genetic correlation (*rg*); 95% highest posterior density (HPD) intervals are given for each value with *l* suffix indicating the lower limit of the 95% interval and *u* suffix indicating the upper limit.

**Supplemental Table 2. Summary of variant-by-sex associations identified in anthropometric traits a)** The number of genetic variants in females and males are listed for the four of the anthropometric traits (waist-hip-ratio, arm-fat-ratio, leg-fat-ratio, and trunk-fat-ratio). **b)** False-discovery rates for traits with more than five variants assigned to a component are across a range of posterior cutoffs (0.5, 0.6, 0.7, 0.8, 0.9). A posterior probability cutoff of 0.8 was used for this paper. The FDR is calculated for each of the male-specific, female-specific, and shared components separately, and indicates the false discovery rate for a variant in that component being associated with that trait. **c)** Model parameter estimates for *pi*, the proportion assigned to each component (null, female, male, and shared respectively, and the *sigmasq* values for the female, male, and shared components are also given (see **Methods**). **d)** Overlap of genes proximal to SEMM identified variant-by-sex associations with literature interactions from [4] [5] (waist-hip-ratio). The table lists the numbers of genes in the reference with sex-specific associations and the number of genes overlapping with these, as well as the identities of these genes.

**Supplemental Table 3. Lists of sex-specific associations in anthropometric traits.** For anthropometric traits, lists of **a)** female-specific, **b)** male-specific, and **c)** shared associations.Trait, variants (ID, chromosome [CHR], position [POS], reference and alternate alleles [REF and ALT], minor allele frequency [MAF]), their associated betas, standard errors, and p-values from GWAS sex-divided summary statistics (*B.f* = beta in females, *B.m* = beta in males) are shown. Posterior probabilities of the variant belonging to the null *(p0)*, female-specific, *(p1)*, male-specific *(p2)*, and shared *(p3)* components. HGNC (GENE column) and variant consequence information (Consequence and HGVSp columns) are also provided.

**Supplemental Table 4. List of Quantitative Traits Examined.** Biomarker traits are listed with their UK Biobank Field ID (UKB ID). Derived phenotypes are from [10].

**Supplemental Table 5. SEMM Genetic Parameter Estimates for Biomarkers**. See the caption for Supplemental Table 1 for a description of table format.

**Supplemental Table 6. Genetic Correlations including Menopausal Status**. The data was divided into pre-menopausal women, post-menopausal women, and men, and SEMM was fit to these data to calculate genetic correlations between pre- and post-menopausal women (*rg.pre_post)*, pre-menopausal women and men (*rg.pre_male)*, and post-menopausal women and men (*rg.post_male*). 95% highest posterior density intervals are given for each value with *l* suffix indicating the lower limit of the 95% interval and *u* suffix indicating the upper limit. The proportion assigned to the non-null component (*pi[1]*) is also provided.

**Supplemental Table 7. Summary of variant-by-sex-effect associations in biomarker traits.** This includes a summary of counts of variants identified per trait **(a)**, FDR estimates (**b**), and model parameter estimates **(c)**. See the caption for Supplemental Table 2 for a longer description of fields included.

**Supplemental Table 8. Sex-specific variants in biomarker traits.** Lists of **a)** female-specific, **b)** male-specific, and **c)** shared variants identified by the SEMM. See the caption for Supplemental Table 3 for a description of the fields.

**Supplemental Table 9. Validation of sex-specific variants in held-out Non-British White cohort.** $R^2$ values are listed for relationship between the discovery (White British) and validation (Non-British White) effect size estimates for the set of sex-specific variants found in the discovery population (either female- specific or male-specific). No values are given for male-specific variants for the anthropometric traits because few or no male-specific variants were identified in these traits.

**Supplemental Table 10. List of traits included in Mendelian Randomization analysis**. Contains the trait, the source of the trait, and the MRBase ID. The table also includes the sample size of the population and whether the population is single sex or both sexes combined.

**Supplemental Table 11. Mendelian Randomization results**. Estimated betas (b), standard errors (se), and p-values (pval) for each trait with female-specific variants as exposures and male-specific variants as exposures. The number of SNPs in the exposures (nsnp) is also shown.

**Supplemental Table 12. Covariate adjustment of testosterone for polygenic risk models**. For each of the three cohorts (male-only cohort, female-only cohort, and the combined cohort of males and females), a linear regression was performed to adjust the covariates. The indicator variable "sex" was used only in the combined cohort. The estimated BETAs (Estimate), its standard errors (Std Err), and the T-values are shown.

### Supplemental Figures

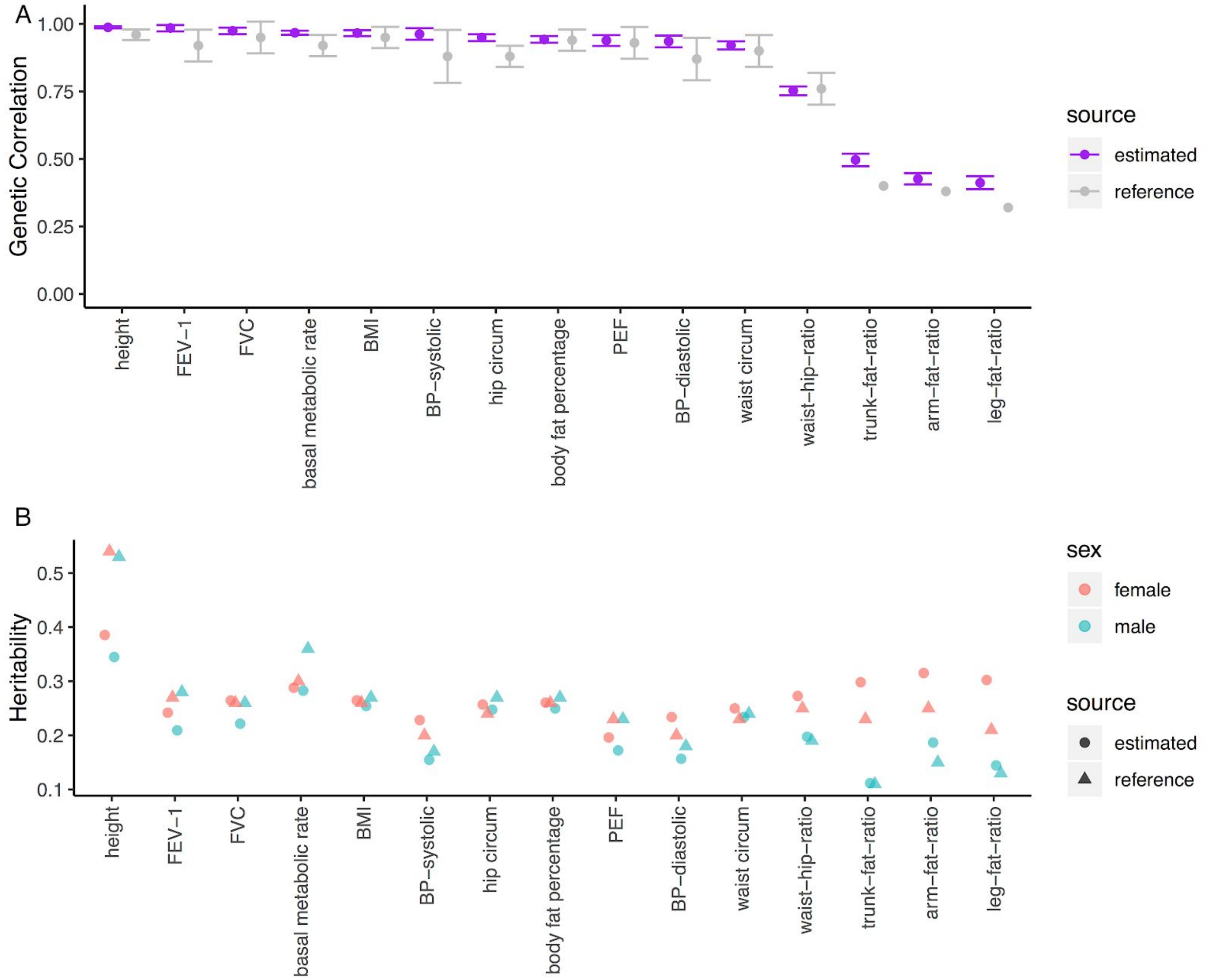


**Supplemental Figure 1. Comparison of estimates for anthropometric traits.** A) Genetic correlation estimates from SEMM (purple) match those previously calculated for a subset of the UK Biobank population (fat-ratio trait estimates from [4], other traits from [22] (gray), 95% HPD interval shown for estimates, 95% CI shown for anthropometric traits where available. B) Heritability estimates (circles) for males (cyan) and females (red) also are similar to those in the literature (triangles).

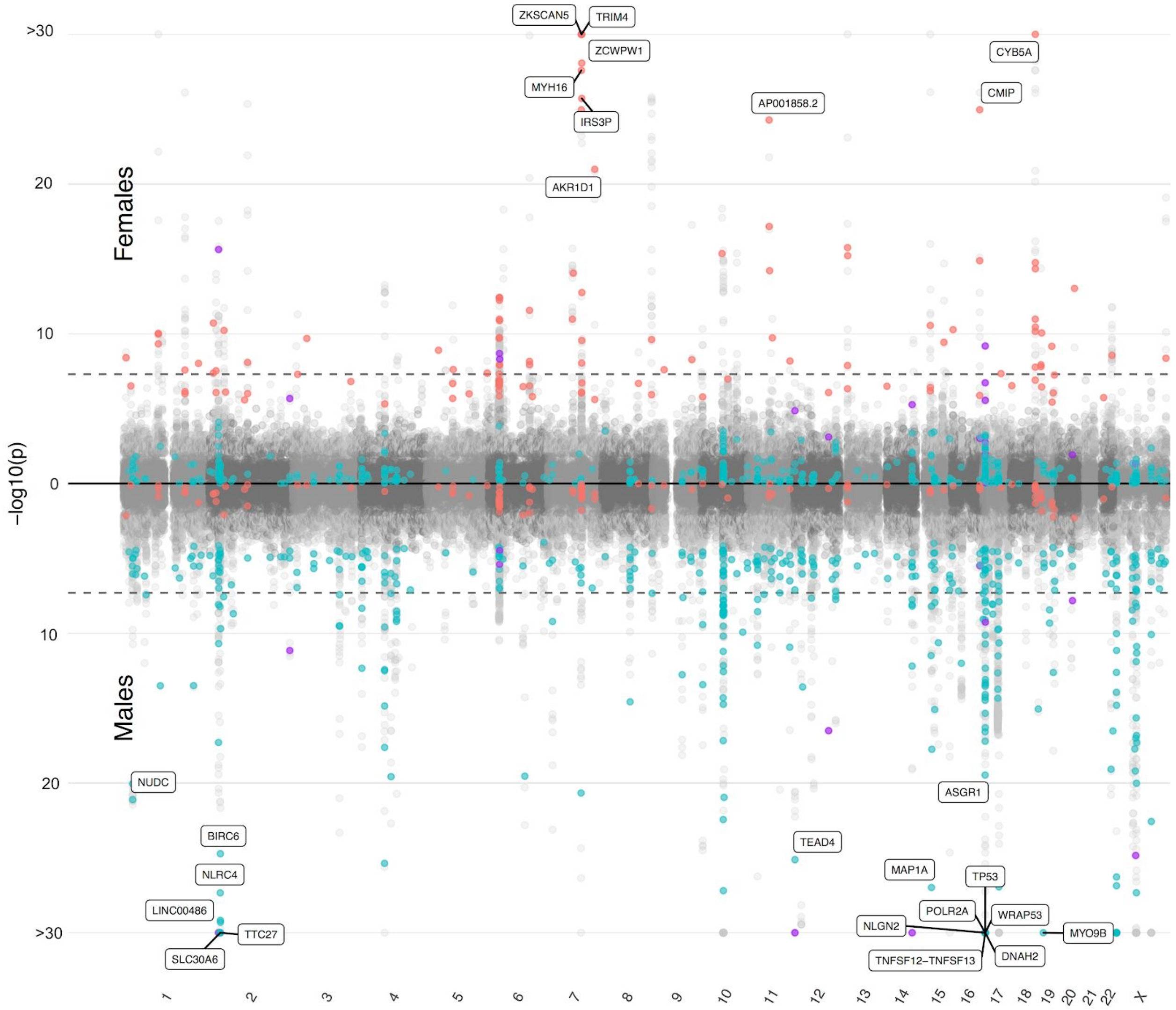


**Supplemental Figure 2. Miami plot showing sex-specific variants associated with testosterone levels.** P-values for females are shown on the top half of the plot, and males are shown on the bottom half of the plot with inverted p-values. Teal dots are male sex-specific variants, red dots are female-specific variants, and purple dots are those assigned to the shared component. Light gray dots are variants that were excluded from the model fitting process because they are in LD with other variants. Each variant is displayed in both the male and female portions of the plot; we can use this to visualize the difference in p-values for the effects. P-values < $10^{-30}$ were truncated to $10^{-30}$ for easier visualization, see **Supplemental Table 8** for the p-values for these variants; and the names of genes proximal to variants with p < $10^{-20}$ in either males or females are shown. **B)** The same plot with missense variants highlighted.

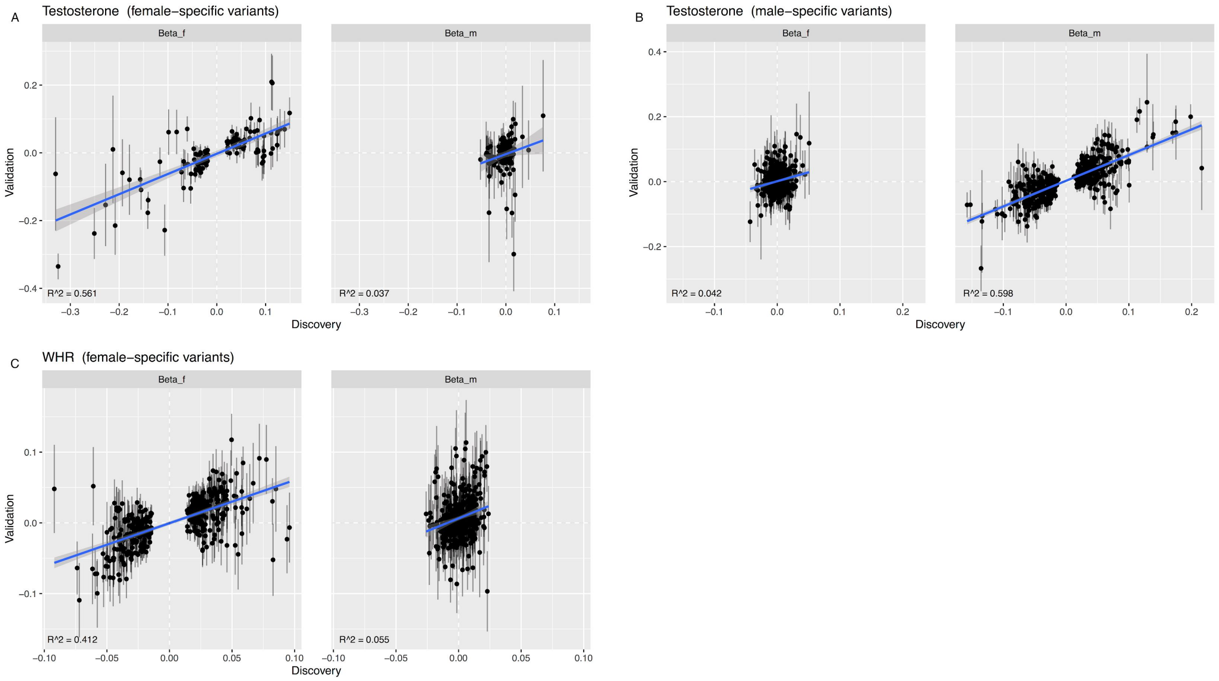


**Supplemental Figure 3. Validation of sex-specific variants in held-out Non-British White cohort.** Each point is the effect size estimate of a variant in the discovery (White British) vs. validation (Non-British White) cohorts, error bars show the standard errors of these estimates from GWAS summary statistics. The estimated effect sizes in females (Beta_f) and males (Beta_m) are given separately. A linear fit is shown in blue with standard error displayed in gray; the $R^2$ associated with this fit is displayed on the bottom left of each figure. Dashed white lines show the no-effect lines (0,0). Female- (A) and male-specific (B) variants in testosterone are shown separately; female-specific variants from WHR are also shown in (C).

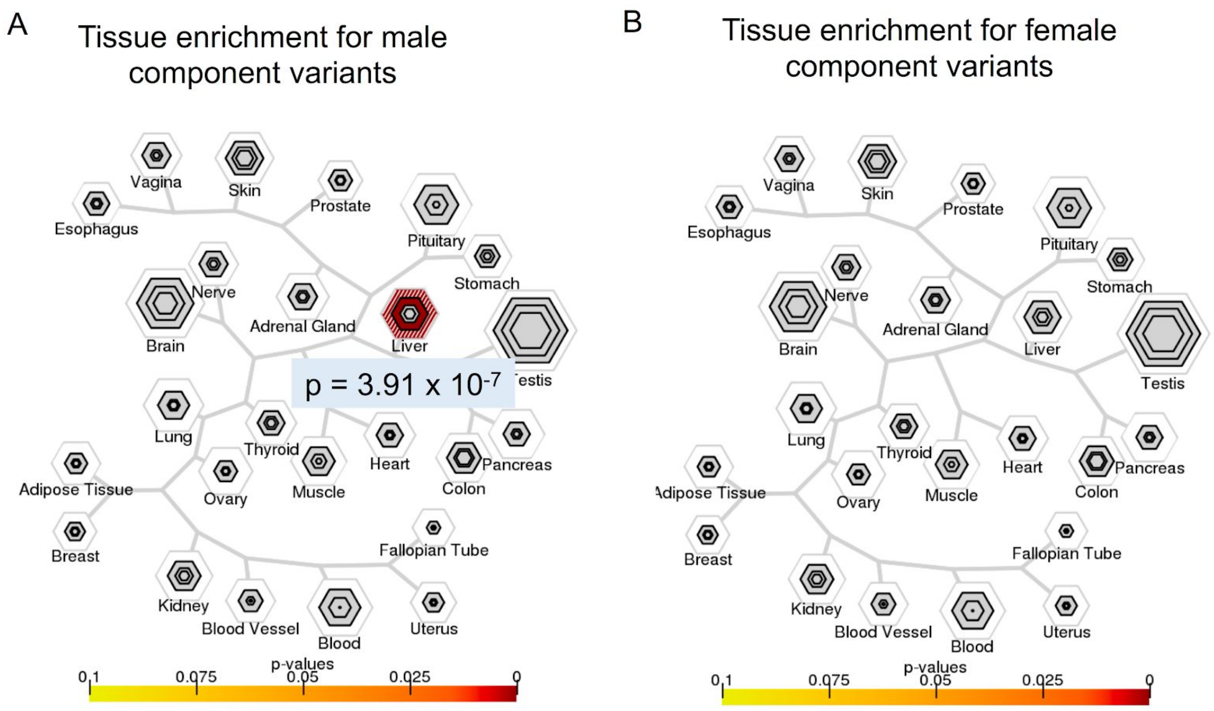


**Supplemental Figure 4. Tissue-specific enrichment analysis for sex-specific genetic effects on testosterone levels.** Visualization of tissue-specific enrichment for male-specific genetic effects on testosterone levels created by the TSEA tool (see Online resource). Each node is a tissue, the color of the tissue colors the enrichment of the set of genes within the genes that are specific to that tissue (as indicated by a Benjamini-Hochberg corrected Fisher's exact test p-value). The size of the node corresponds to the number of enriched transcripts, each node contains three subsections, these represent three different stringency thresholds for the selection of tissue-specific genes. The topography of the diagram is generated from hierarchical clustering of the transcripts in each tissue, nodes closer to each other share more transcripts [62]. Liver is highlighted with significant enrichment in males **(A)** ($p = 3.91x10^{-7}$) and but not females **(B)** ($p > 0.1$).

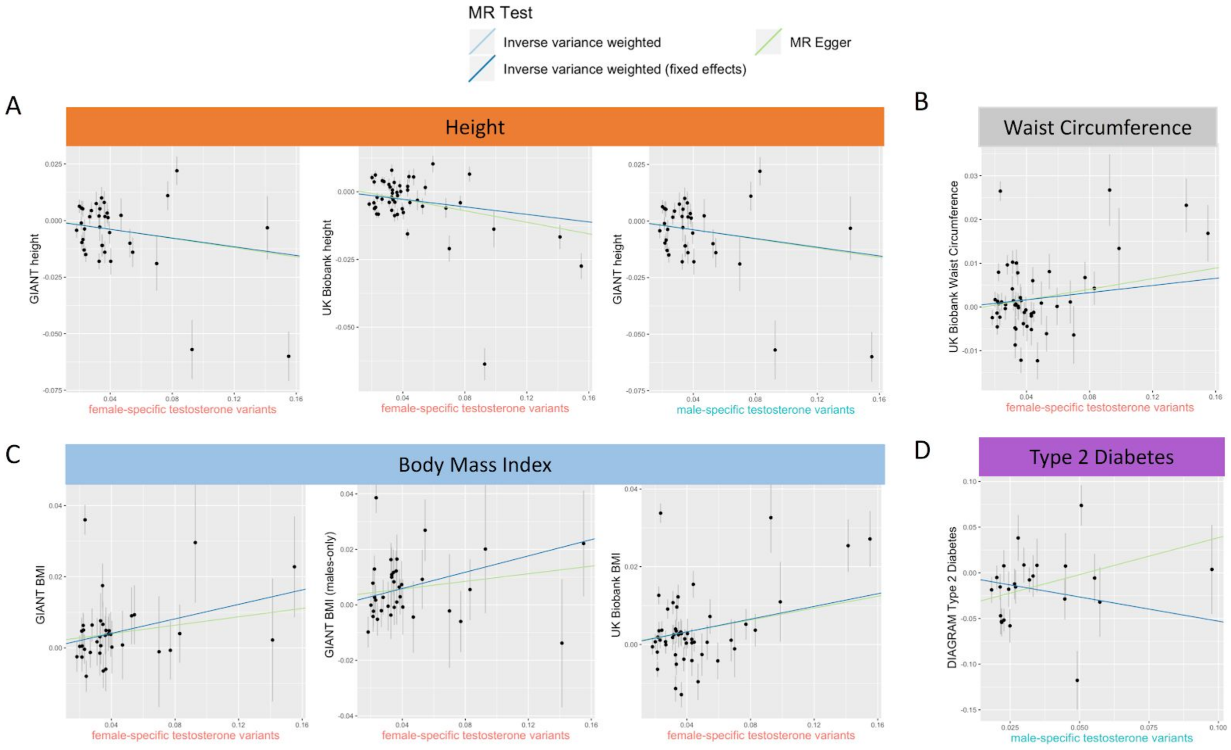


**Supplemental Figure 5. Mendelian Randomization results where sex-specific testosterone variants show evidence of a causal effect on the trait.** Each plot shows the SNP effects for one set of variants (either male or female variants) and one outcome or trait. Mendelian Randomization fits are shown for Inverse variance weighted (light blue), inverse variance weighted fixed effects (blue), and MR Egger (light green). Female-specific variants show evidence of a causal association with height (A), waist circumference (B), and body mass index (BMI) (C); male-specific variants show evidence of causal associations with height and Type 2 Diabetes (D) (DIAGRAMplusMetaboChip). For height and BMI, both outcomes in from the UK Biobank (far left A and C) and GIANT consortia show associations, and for BMI GIANT data both male (center C) and combined (right C) populations show this relationship.

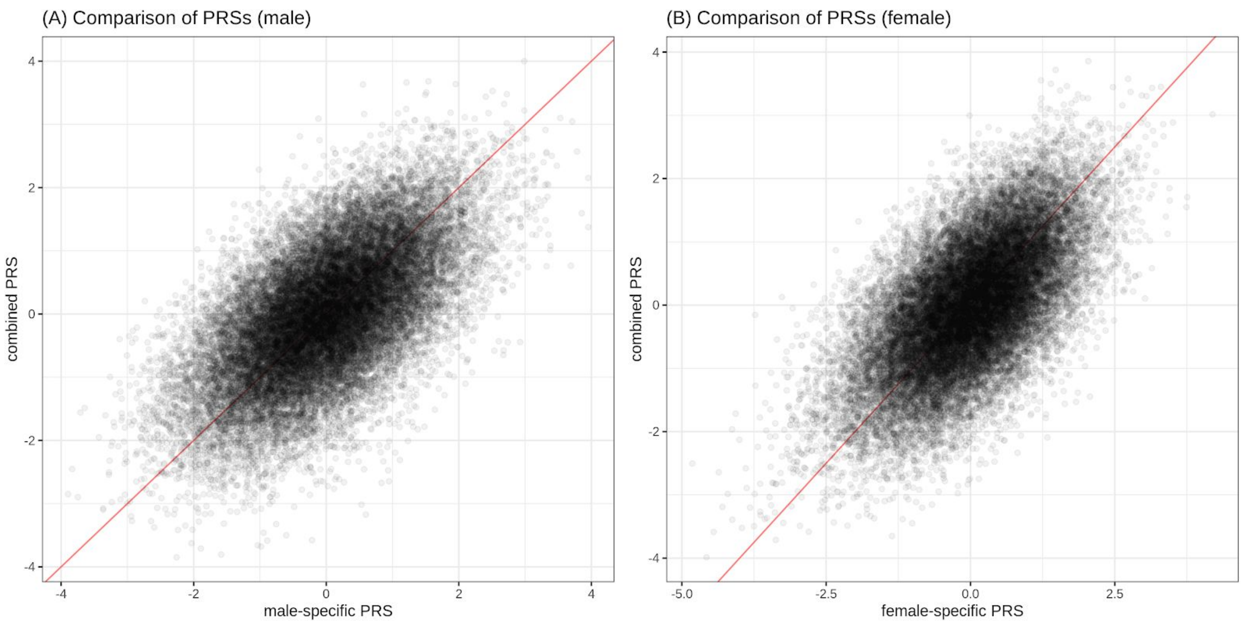


**Supplemental Figure 6. Comparison of sex-specific PRS and the combined PRS.** The sex-specific PRS (x-axis) and the combined PRS (y-axis) are shown for the individuals in the test set for males (A) and females (B). The two PRSs are centered and scaled so that each of them has zero mean and unit variance. The red diagonal line indicates y = x.
